# Predicting the Patterns of Kinship Dynamics in Human Societies

**DOI:** 10.1101/2025.04.13.648649

**Authors:** Peng He, Mark Dyble, Rufus A. Johnstone

## Abstract

By shaping the inclusive fitness effects of behaviours, kinship environments have important implications for social evolution, and studies across non-human animal societies have shown that the patterns of kinship dynamics individuals experience as they age can be predicted by simple demographic parameters relating to sex-specific dispersal patterns, the extent of extra-group mating, and the age-specific patterns of mortality and fertility. However, it remains unclear whether these insights apply within our own species, where we see both extended lifespans in which kinship dynamics can play out, and remarkable variations in cultural practices relating to marriage and residence that influence the patterns of inter-individual relatedness. Here, using a newly developed model of kinship dynamics, we explore how marriage practices and post-marital residence rules, along with the typical survival and fertility schedules of our own species, can predict the emergent patterns of female and male kinship dynamics in human societies. By showing evidence that the model successfully captures the patterns of female and male kinship dynamics underlying human societies, we demonstrate that, despite the diversity in human social organisation, a relatively simple set of parameters relating to male and female rates of dispersal and the prevalence of extra-group mating suffices to capture the qualitative patterns of human kinship dynamics. Given the inclusive fitness implications kinship dynamics have for sex- and age-linked variations in behaviours, our study thus generates new insights into human social evolution.

## INTRODUCTION

Social behaviours vary significantly across mammals, from largely solitary species to those living in cooperative breeding groups^1,2^. A key driver of such variation is the degree of genetic relatedness between groupmates — in line with kin selection theory^3-5^, higher levels of genetic relatedness among groupmates are generally associated with higher frequencies of cooperative behaviours such as alloparental provisioning^6,7^ and divisions of labour^8,9^. While such associations between the ‘static’ views of both inter-individual relatedness and the expression of social behaviours are widely acknowledged, recent empirical and theoretical studies across non-human social animals have also shown that an individual’s average relatedness to its groupmates can vary systematically with sex and age following processes such as birth/death, immigration/emigration, and mating in groups^10^, and such predictable variations can potentially drive sex- and age-linked trends in social behaviours^11,12^. Across the sexes, for example, in species in which males disperse upon sexual maturity while females stay in their natal group, adult females are, on average, more closely related to their groupmates than males are^11^. Such sex differences have been shown in species including lions^13^, badgers^14^, and Rhesus macaques^11^ and may drive important sex differences in social behaviour^10,11^. Individuals may also vary in their average relatedness to their group as they age. For example, female killer whales are increasingly related to their groups as they are reproductively active but become increasingly unrelated to their group after reproductive cessation^15,16^. Taken together, such sex- and age-linked variations in individuals’ average relatedness to their social group, termed as kinship dynamics^10^, may drive notable variations in social behaviours within and between species, and may produce evolutionary conflicts-of-interest between sexes^17-19^ and generations^15,20,21^.

Despite these cross-species insights into the emergent patterns of kinship dynamics and their inclusive fitness implications for social behaviours, we know little about how individuals’ average relatedness to their group varies with sex and age within our own species. For humans, any attempt at characterising such species-wide patterns of kinship dynamics is obscured by the considerable cross-cultural diversity seen in our social organisation. Human societies vary substantially in dispersal (or residence) rules, marriage practices, and social structures^22-26^. In dispersal, for example, we see patrilocal residence in which women disperse upon marriage, matrilocal residence in which males disperse on marriage, multilocal residence in which a household moves frequently between sites, duolocal residence in which neither sex disperses, and neolocal residence where new couples move to a new location entirely^22,27^. In marriage, we see examples of monogamy, polygyny, and polyandry^19,24,28^ as well as variable rates of marriage dissolution^29^ and extra-pair paternity^30^. Where this variation creates age and sex differences in individuals’ average relatedness to their households, theory would suggest that this may result in age and sex inequality in political, economic, and social life^27,31-33^. Some recent studies support this possibility. For example, recent studies have demonstrated that women living with their own kin (rather than their husband’s kin) have an early age at first birth^34^ and greater bargaining power in farm labour, working less hard as a result^27^. We suggest that consideration of the causes and consequences of cross-cultural variation in human kinship dynamics can offer important insights into variation in sex and inter-generational conflicts of interest across societies.

Here, we use a newly developed theoretical model to explore the consequences of sex- and age-linked variations in dispersal, fertility, and mortality for the emergent patterns of human kinship dynamics, and test our predictions with 11 cross-cultural empirical datasets across the global. We demonstrate how the patterns of female and male kinship dynamics in human societies can be predicted by simple sets of characteristic demographic parameters, and discuss the inclusive fitness implications of our predictions for human social behaviours — such as invoking local relatedness asymmetries between the sexes to explain cultural variations in inter-generational transfers of socio-political authorities across societies.

## METHODS

For clarity and simplicity, we focus on the patterns of kinship dynamics both sexes experience in the typical (**i**) bilocal society (BIL), where the dispersal of a female is as likely as that of a male and all the locally born offspring in a group are sired by the males from the same group, (**ii**) duolocal society (DUO), where none of the sexes disperse and all the locally born offspring in a group are sired by non-local males (i.e., those from elsewhere), (**iii**) matrilocal society (MAT), where only males disperse, and all the locally born offspring in a group are sired by the males from the same group, and (**iv**) patrilocal society (PAT), where only females disperse and all the locally born offspring in a group are sired by the males from the same group. These typical types of human societies are well acknowledged in anthropology, and the clear-cut of their definitions and realizations allows us to closely test our model predictions with empirical data available. Below, we first generate predictions for these societies with a newly-developed model of kinship dynamics^16^, and then test our predictions with published relatedness datasets available across these four type of human societies.

### The model of kinship dynamics

We generate the patterns of female and male kinship dynamics across the above-mentioned societies with the ‘hybrid’ version of the model described by He et al.^16^, which incorporates the effects of age-related *changes* in survival and fertility of the sexes on individuals’ average relatedness to their groups, while considering the effects of dispersal and mating as its predecessors do^11,12^. Briefly, the model considers an infinite, diploid, bisexual population, divided into discrete groups — with given numbers of females and males in each group. As discrete timesteps proceed, the age compositions in a group change following given sex-age-specific survival rates, and such transitions of group age compositions are captured by a Markov process. It assumes female population demographic dominance, where the number of offspring produced depends only on the age-specific fertilities of females (the age-specific fertility of a male is the amount of paternity he enjoys over all the offspring reproduced in the population — by competing with other males under the given rate of local/non-local mating). The number of offspring produced in the population is assumed large enough so all the breeding vacancies in a group at any timestep are fully occupied (but females vary in their *relative* fertility — described by age-specific fertilities). At each timestep a given proportion of the offspring produced by each female are females while the rest are males, and a given proportion of all the offspring produced in a group are sired by local males while the rest by non-local males. An offspring disperses to a random group with its sex-specific dispersal rate. After dispersal, offspring in each group (natives and immigrants) compete in a ‘fair lottery’ for the breeding vacancies created by the deaths of others of their own sex in the group — those failed to obtain any vacancy die, and these demographic cycles then repeats. Kinship dynamics are derived as individuals’ sex- and age-specific average relatedness to their groupmates at population demographic equilibrium (where relatedness between individuals is calculated as the probability of *identity by descent* of two homologous neutral genes, each of which is randomly sampled from the same loci in each individual; see in He et al.^16^).

### Model parameters for predictions

In terms of model parameters, the four types of human societies are characterized by the combinations of (**1**) the rates of female and male dispersal (i.e., *d*_*f*_ and *d*_*m*_, respectively) and (**2**) the rate of local vs non-local mating (i.e., *m*, the probability that an offspring born in a group is sired by local versus non-local males), such that the BIL, DUO, MAT and PAT societies are explicitly captured by the parameter sets {*d*_*f*_ = .5, *d*_*m*_ = .5, *m* = 1}, {*d*_*f*_ = 0, *d*_*m*_ = 0, *m* = 0}, {*d*_*f*_ = 0, *d*_*m*_ = 1, *m* = 1} and {*d*_*f*_ = 1, *d*_*m*_ = 0, *m* = 1}, respectively. For simplicity and the robustness of predictions, we assume the sex- and age-specific survival rates and fertilities are consistent across the four types of human societies; that is, when generating predictions, we consider the effects of the survival and fertility schedules that are common to our own species across individual societies. For generality, we use the 1986 global human survival and fertility schedules published by the UN^35^ as model inputs. Due to the unavailability of male fertility data, we assume males share the same fertility curve as females (Fig. A1 in the appendix). As group (or household) sizes are typically fluid and can vary across human societies (and can modulate the predicted patterns of kinship dynamics^36^), we generate predictions over varying group sizes and sex compositions, which are derived by the unique combinations of the numbers of females and males from the set {2, 3, 4, 5}, such that group sizes range from 4 to 10 while sex ratios range from 2: 5 to 5: 2.

### Testing model predictions with relatedness data across human societies

We test model predictions with the empirically observed patterns of female and male kinship dynamics across the four typical types of 11 human societies, as well as the age-specific relatedness asymmetries between the sexes to their group based on these observed patterns of kinship dynamics. Specifically, we use the relatedness data empirically collected from (**i**) the *bilocal* (BIL) *Agta* society (*n* = 1, labelled as ‘AG’) described by Dyble et al^37^, (**ii**) the *duolocal* (DUO) *Mosuo* society (*n* = 1, labelled as ‘MS’) described by Wu et al^38^, (**iii**) the *matrilocal* (MAT) *Pumé* and *Mayangna* societies (*n* = 2, labelled as ‘PU’ and ‘NI’, respectively) and (**iv**) the *patrilocal* (PAT) *Maya, Lamalera, Gambia, Mosuo, Alakāpuram, Tenpaṭṭi*, and *Tanna* societies (*n* = 7, labelled as ‘MA’, ‘LA’, ‘GA’, ‘MP’, ‘AZ’, ‘TP’, and ‘TA’, respectively) described by Koster et al^25^ (Table A1 in the appendix). For each of these empirical societies, we calculate for the sex- and age-specific average relatedness of individuals to their groups, and use these sex- and age-specific values to identify the ‘best-fit’ predictions from those candidates generated with varying group sizes and sex compositions (described above), such that the corresponding ‘best-fit’ numbers of females and males minimize the sum of the squared deviations of the predicted relatedness values (across both sexes and all ages) from those observed (across both sexes and ages).

With the ‘best-fit’ relatedness predictions for the sexes (summarized in Fig. 1-A below) for each of the empirical society, we statistically test whether the model successfully captures the systematic patterns of female and male kinship dynamics underlying the four typical types of human societies, in terms of both (**1**) the sex- and age-specific average relatedness values and (**2**) the age-specific natural log ratios of *female* to *male* mean unrelatedness values (i.e., 1 − mean relatedness), as a measure of the age-specific average relatedness asymmetry between the sexes to their groups — the more the positive/negative log ratios deviate from 0 (i.e., when individuals of both sexes at a given age are *symmetrically* unrelated/related to their group), the more the males/females are related to their group than their peer females/males are. To do this, we construct a test statistic *sMSD*, i.e., the sum of the mean squared deviations of the set of 11 empirical observations from the associated set of 11 ‘best-fit’ relatedness predictions. Specifically, in terms of sex- and age-specific average relatedness, the mean squared deviation (*MSD*) of the observations from the associated particular ‘best-fit’ predictions for *each* empirical society is derived across the sexes and all the ages present in the dataset for the empirical society; in terms of age-specific natural log ratios of female to male mean unrelatedness, the mean squared deviation of the observations from the associated particular ‘best-fit’ predictions for *each* empirical society is derived across all the ages shared by both sexes in the dataset for the empirical society.

**Fig. 1.**
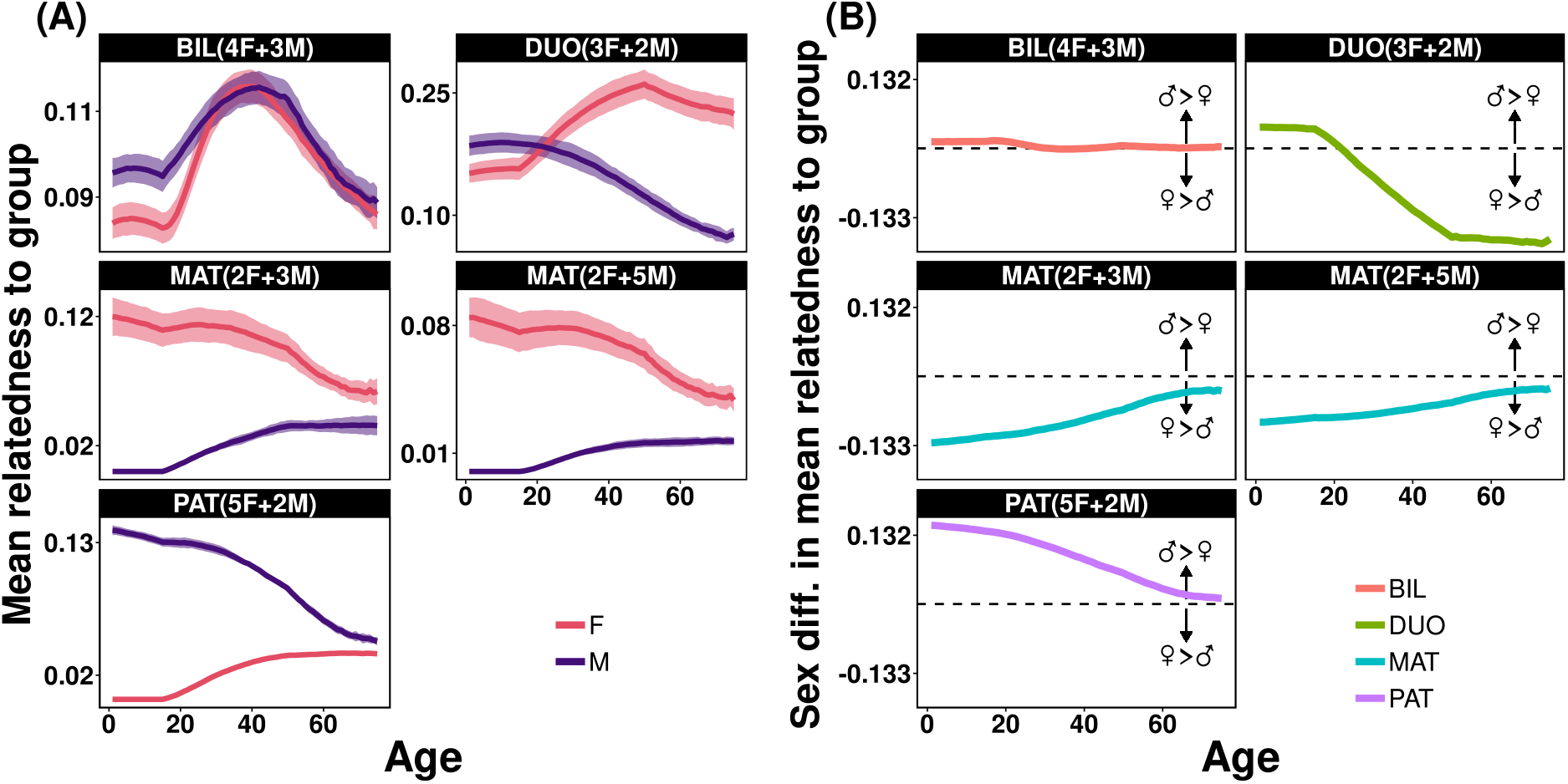
(**A**) The summary of the predicted patterns of female (F, red) and male (M, purple) kinship dynamics in the *bilocal* (BIL), *duolocal* (DUO), *matrilocal* (MAT) and *patrilocal* (PAT) societies (i.e., each panel) that best explain those empirically observed in each of the 11 human societies across these society types (shaded ribbons indicating the range of *mean* ± sd from 10 replications with the ‘hybrid’ version of the model of kinship dynamics described in He et al^16^), and (**B**) the summary of the age-specific (natural) log ratio of female to male mean *unrelatedness* (1 − mean relatedness) derived from those predictions in (A), which measures age-linked relatedness asymmetry between the sexes — the more the positive (or negative) values deviate from the horizontal dashed black lines of symmetry, the more the males (or females) are related to their household (or group) than their peer females (or males) are (biases towards the sexes are indicated by the arrows in the panels). The abbreviations for the four typical societies in question are given in the black strip in each panel, along with those ‘best-fit’ numbers of females (F) and males (M) assumed in each household (or group) by the model for each of the 11 human societies, such that they minimise the sum of squared deviations of predicted mean relatedness values from those empirically observed in individuals across the sexes and all ages in each of the societies (see methods).

We then use both the permutation- and the randomization-based approaches to derive the reference distributions for the *sMSD*, and evaluate the statistical significance of the observed test statistic (which is derived by associating the observations from each of the 11 societies with the corresponding ‘best-fit’ predictions for that society). To derive the distributions, we calculate the resulting test statistic *sMSD* while randomly associating the set of 11 ‘best-fit’ predictions with those 11 empirical observations across societies, by means of either permutating or bootstrap sampling those 11 ‘best-fit’ predictions. When constructing the reference distributions, we use *sMSD* derived from those unique permutations for the full set of 11 ‘best-fit’ predictions for the empirical societies, and set bootstrap sample size the same as the number of unique permutations, while keeping the probability that a given ‘best-fit’ prediction being bootstrap-sampled in each trail the same as its relative frequency in the whole set of 11 ‘best-fit’ predictions for the empirical societies. With the 11 empirical relatedness dataset, we identified a set of 6 unique predictions across the four typical types of human societies (Fig. 1) — where (**i**) the *bilocal* AG is associated with the prediction with 4 females and 3 males, (**ii**) the *duolocal* MS is associated with the prediction with 3 females and 2 males, (**iii**) the *matrilocal* PU and NI are associated with the predictions with 2 females and 3 males, 2 females and 5 males, respectively, while (**iv**) the *patrilocal* MA, LA, GA, MP, AZ, TP, and TA are consistently associated with the prediction with 5 females and 2 males (Fig. 1). With these ‘best-fit’ predictions for the 11 empirical societies, we derived 11!/ (7! 1! 1! 1! 1!) = 7,920 unique permutations of them. Based on the derived distributions of *sMSD*, we compute their critical (left tail) quantiles of statistical significance for testing the predicted patterns of kinship dynamics and their sex differences across age (with significance level of 0.05 across all tests).

## RESULTS

### Predicted patterns of kinship dynamics and their sex difference

In the bilocal (*BIL*) society, both female and male average relatedness to their group exhibit a ‘three-stage’ pattern: it decreases (though slightly) with age in childhood, then it increases once individuals reaching sexual maturity (with female’s rate of increases higher than that of males’), and it decreases again from midlives (Fig. 1-A). These sex-specific patterns of kinship dynamics lead to similar local kinship environments both sexes experience over age (Fig. 1-B). In the duolocal (*DUO*) society, female kinship dynamics also exhibit a ‘three-stage’ pattern as in the bilocal society, but girls’ average relatedness to their group tend to increase (though slightly) across their childhoods (Fig. 1-A). In such societies, males tend to be increasingly less related to their group as they age (Fig. 1-A). These sex-specific patterns of kinship dynamics give rise to notable age-linked relatedness asymmetry between the sexes — females initially are less related to their group when compared with their peer males and then tend to be increasingly more related to their group than males do afterwards (and remain so in their late lives). The patterns of kinship dynamics the sexes experience in the matrilocal (*MAT*) society exhibit the opposite trend of each other, where females are increasingly less related to their group over age while males are increasingly more related to their group as they age (Fig. 1-A). These opposite trends of kinship dynamics lead to the long-lasting relatedness asymmetries between the sexes — females are more related to their group when compared to their peer males, though they are less so as they age (Fig. 1-B). In the patrilocal (*PAT*) society, the patterns of female and male kinship dynamics also exhibit the opposite trend of each other as in the matrilocal society, but females instead are increasingly related to their group as they age while males are increasingly less related to their group over age (Fig. 1-A), and these patterns of kinship dynamics also lead to the consistent relatedness asymmetries between the sexes — in clear contrast to that in the matrilocal society, males in the patrilocal society are more related to their group than their peer females, though they are less so as they age (Fig. 1-B).

### Test model predictions with empirical datasets

While empirical datasets reveal notable fine-scale variations in individuals’ average relatedness to their groupmates across age (Fig. A2), there are also qualitatively identifiable patterns of female and male kinship dynamics across each of the four typical types of societies that fit model predictions. For example, in the bilocal AG society^37^, the average relatedness of individuals of both sexes to their group decreases with age while juvenile, and then it increases before decreasing again in midlife (Fig. A2), and the local kinship environments both sexes experience tend to remain consistent over age (Fig. 2); in the duolocal MS society^38^, females are increasingly more related to their group than their peer males are as they age (Fig. 2), while in the matrilocal PU and NI societies^25^, females are likely to be more related to their group than their peer males are (Fig. 2).

**Fig. 2.**
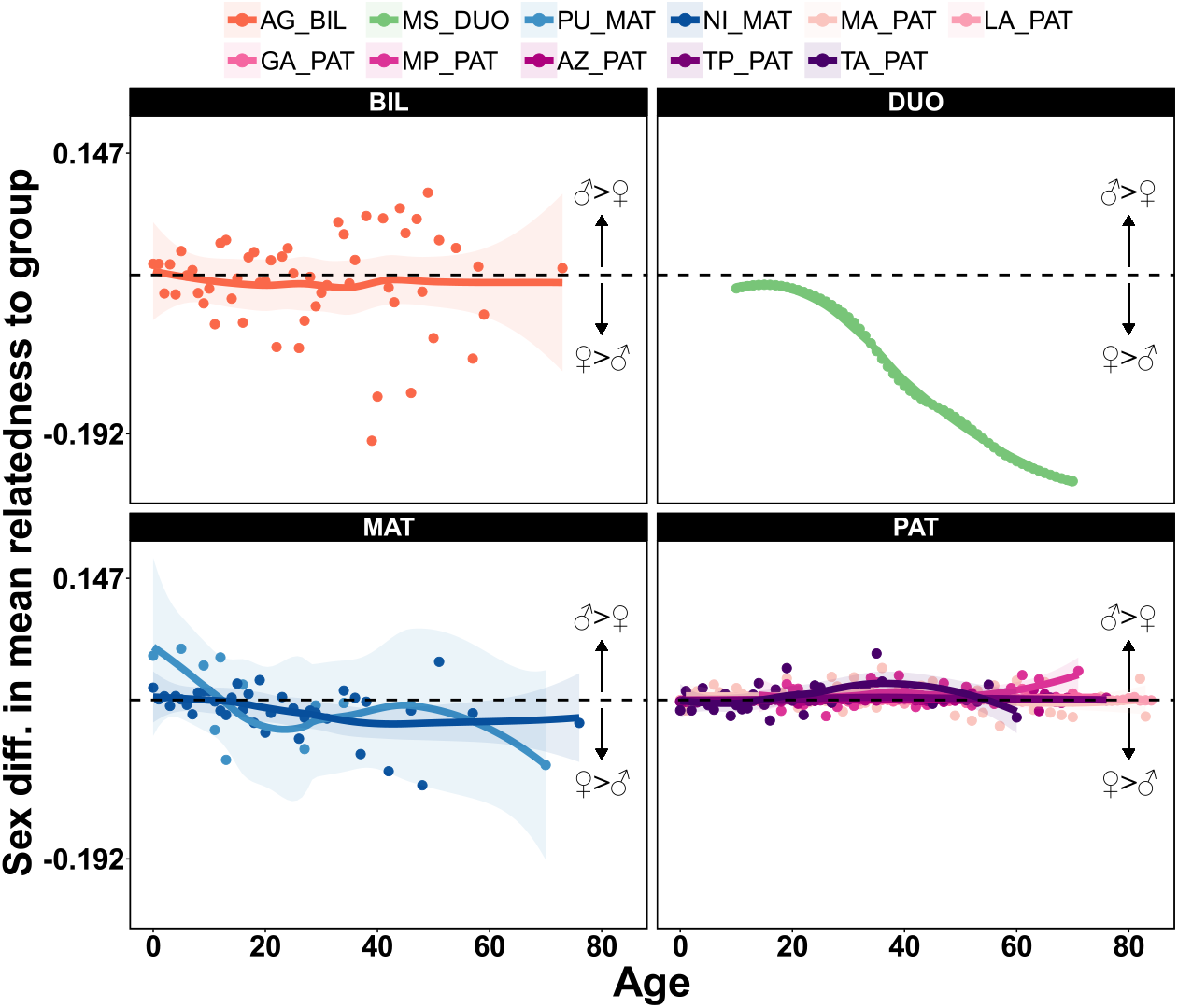
The empirically observed patterns of age-specific relatedness asymmetries between the sexes to their group (i.e., the age-specific natural log ratio of female to male mean unrelatedness as described earlier) across the 11 human societies, each of which is classified as either the *bilocal* (i.e., BIL, prefixed by ‘AG’ as the abbreviation for the Agta society^37^ in the legend), *duolocal* (i.e., DUO, prefixed by ‘MS’ as the abbreviation for the Mosuo society^38^ in the legend), *matrilocal* (i.e., MAT, prefixed by ‘PU’ and ‘NI’ as the abbreviations for the *Pumé* and the *Mayangna* societies^25^ in the legend, respectively), or *patrilocal* (i.e., PAT, prefixed by ‘MA’, ‘LA’, ‘GA’, ‘MP’, ‘AZ’, ‘TP’ and ‘TA’ as the abbreviations for the *Maya, Lamalera*, one of the *Gambia*, the patrilocal *Mosuo, Alakāpuram, Tenpaṭṭi*, and the *Tanna* societies^25^ in the legend, respectively). Filled circles indicate the age-specific observations across the 11 societies while solid lines visually delineate the estimated local trends of these observations over ages (i.e., by the LOESS regression, where the shaded ribbons indicate the 95% confidence intervals), with each colour indicating an empirical human society. The more the positive/negative values deviate from the horizontal dashed black lines of symmetry, the more the males/females are related to their household (or group) than their peer females/males are (biases towards the sexes are indicated by the arrows).

Both the permutation- and randomization-based statistical test results suggest that the model successfully captured the systematic patterns of female and male kinship dynamics underlying the four typical types of human societies (Fig. 3). Specifically, in terms of individuals’ average relatedness to their groups across sexes and ages, the sum of the mean squared deviations (*sMSD*) of the set of 11 empirical observations from the associated set of 11 ‘best-fit’ predictions (summarized in Fig. 1-A) are significantly lower than that would be expected if the set of 11 empirical observations are randomly associated with the set of 11 ‘best-fit’ predictions, by means of either permutating (‘per’ in Fig. 3-A) or bootstrap sampling (‘rnd’ in Fig. 3-A) the set of ‘best-fit’ predictions. Likewise, in terms of individuals’ age-specific (natural) log ratio of mean unrelatedness to their groups, the *sMSD* of the set of empirical observations (across the 11 human societies) from the associated set of 11 ‘best-fit’ predictions (summarized in Fig. 1-B) are significantly lower than that would be expected if these observations are randomly associated with the set of 11 predictions, by means of either permutating (‘per’ in Fig. 3-B) or bootstrap sampling (‘rnd’ in Fig. 3-B) these predictions.

**Fig. 3.**
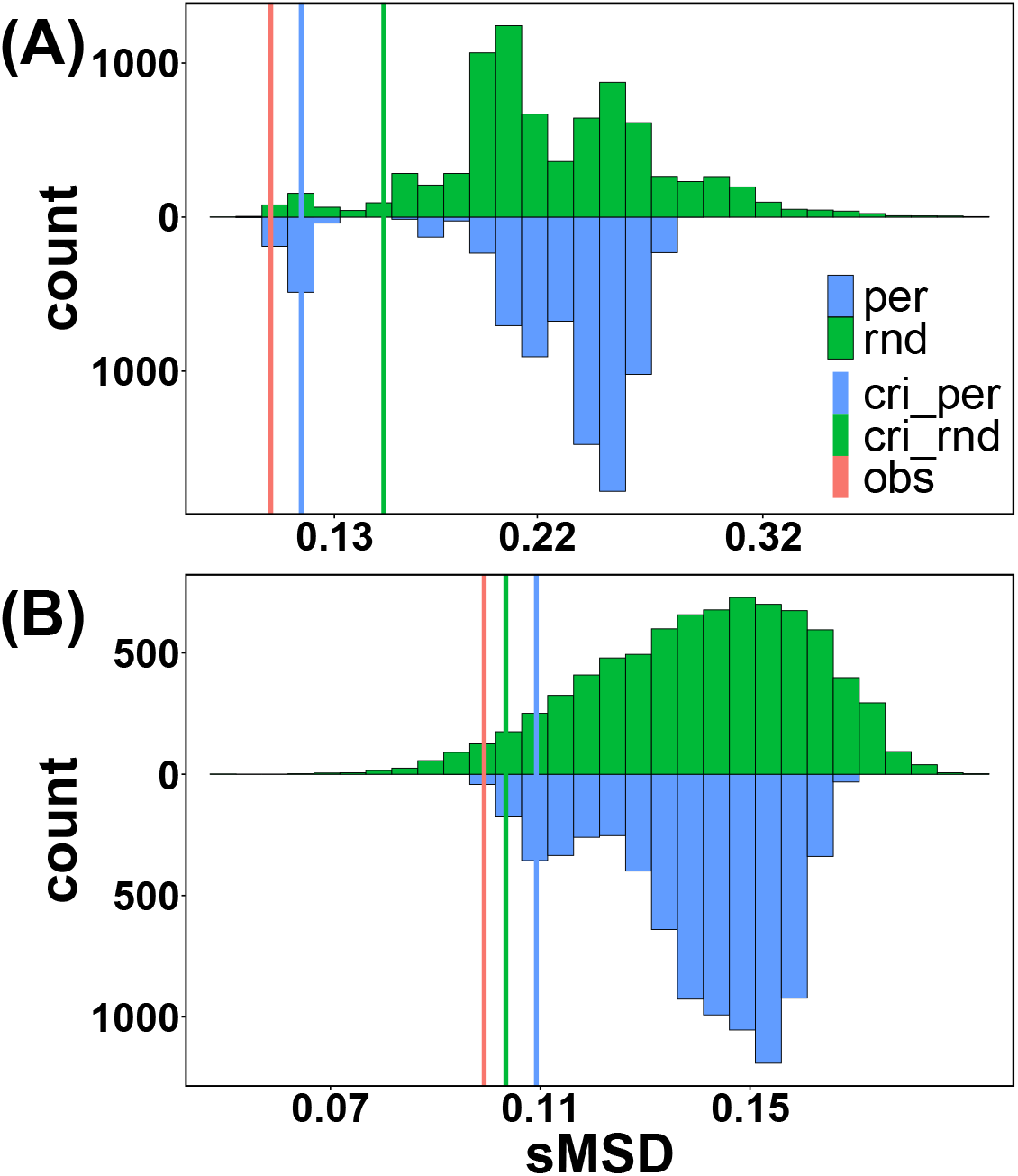
The results of the permutation- and randomization-based statistical tests for model predictions with the empirically observed (**A**) patterns of female and male kinship dynamics and (**B**) patterns of sex differences in kinship dynamics across the 11 human societies. The observed test statistic, i.e., the red vertical lines (labelled ‘*obs*’), is the sum of the mean squared deviations (*sMSD*) of the set of 11 empirical observations from the associated set of 11 ‘best-fit’ predictions. The distributions of the *sMSD* are derived by calculating the resulting test statistic *sMSD* while randomly associating the set of ‘best-fit’ predictions with those empirical observations across the 11 human societies, by means of either permutating (‘per’ and blue) or bootstrap sampling (‘rnd’ and green) those 11 ‘best-fit’ predictions for each of the empirical societies. The blue (labelled ‘cri_per’) and green (labelled ‘cri_rnd’) vertical lines mark the critical (lower tail) quantiles of statistical significance for the tests with the distributions derived with permutations and randomizations, respectively (with the given significance level of 0.05).

## DISCUSSION

Human social behaviours are shaped by kinship ties^23,26,31,33^, and studies across non-human species have demonstrated how patterns of kinship dynamics both sexes experience are linked to fundamental demographic processes in social groups^10-12,16^. Using a recently developed model and empirical relatedness datasets available, we demonstrate, for the first time, how the qualitative patterns of sex- and age-link variations in individuals’ average relatedness to their group can be predicted by simple demographic characteristics of human societies. By doing so, our study generates new insights into sex- and age-linked trends in human social behaviours.

By showing how marriage practices and post-marital residence rules are linked to the dynamic kinship environments individuals of both sexes experience over age, our study offers biologically sensible explanations for variations in human social behaviours. In anthropology, how matrilineal social systems (which are associated with matrilocal residence) cope with the seemingly ‘split-up’ of the role of a man as the husband/father versus brother/uncle of a woman/child has intrigued researchers for a long time as ‘the matrilineal puzzle’^39^, which arises from the inconsistency where descent is typically through women but authorities (over resources etc.) are often vested in men^40^, whereas both descent and inheritance are men-centralized in the patrilineal societies (which are associated with patrilocal residence). We note that while an in-marrying man is increasingly related to his group as he ages in the matrilocal societies, he may still be less related to his own than to his natal group — as we may expect from the low relatedness plateau he eventually reaches in late life. Such a slow build-up of a man’ kinship ties to his own household over age, along with the low paternity confidence he may experience^41^, help to explain why men tend to predominantly invest resources in their matrilineal sisters’ instead of their own sons^42,43^. What remains to be tested based on our predictions is, however, whether such age-linked increases in relatedness of an in-marrying man to his own group are positively correlated with his investments in the group as he ages.

The patterns of age-linked relatedness asymmetries between the sexes to their group may also help to generate deeper insights into sex difference in behaviours^44,45^ and gender inequality in social, economic, and political life in human societies^27,32,46,47^. Across human societies, studies have shown that the social behaviours of the sexes are linked to residence rules. For example, in hunter-gatherer societies, males tend to contribute less to subsistence in patrilocal but more in matrilocal societies than females do^48^; in agriculturalists and pastoralists, females tend to reproduce earlier when co-residing with their own parents than with non-kin^34^, and their higher workload (relative to males) is even more pronounced in patrilocal than duolocal societies, while such imbalance tends to be absent in matrilocal societies^27^. As we show, sex-biased dispersal creates systematic age-linked relatedness asymmetries between the sexes to their group. Such asymmetries can generate variations in how much kin-directed support or help (e.g., bargaining power, transfer of resources) is vested between the sexes, and thus explain sex differences in social behaviours^27,34^. Yet, our results stimulate predictions on age-linked trend in sex differences in social behaviours. For example, as both sexes tend to be less asymmetrically related to their group as they age in the matrilocal and patrilocal societies, we may expect less pronounced sex differences in social behaviours in later than earlier life stages, in contrast to those in the duolocal society where more striking sex differences may be expected in later life. We suggest that, along with sex difference in fundamental biology, age-linked variations in the kin environments the sexes experience may also drive sex-differentiated behaviours^44,45^ and gender inequality^46,49^.

In summary, while a society may not be characterized exactly by any of the idealized patterns of mating and post-marital residence as we theoretically define and explore, we suggest that the more it approximates any of these types, the more the observed patterns of human kinship dynamics will fit our predictions, under the general effects of mating and residence patterns on local genetic structures^50-52^. Similarly, while the demographic schedules observed from a given society may deviate from those we explored, we suggest that the predicted patterns of kinship dynamics qualitatively still hold, as these schedules are underpinned by the fundamental biology typical of our own species, such as more rapid increases in mortality in later than earlier adult lives for both sexes^53^ and a significantly prolonged female post-reproductive lifespan^54^. We thus highlight the new insights our study generates into our understanding of human kinship dynamics and their evolutionary implications for sex- and age-linked trends in human social behaviours.

## DATA & CODE AVAILABILITY

All the data and code used in this study are openly available at *https://github.com/ecopeng/Kinship_Dynamics_Human_Societies*.

## ACKNOWLEDGEMENTS

We thank the authors for making their empirical data openly available to allow us to test our predictions.

## AUTHORS’ CONTRIBUTIONS

All authors contributed to the conceptualization, data analyses and writing for this study, and gave final approval for the publication of the study.

## REFERENCES

1 Kappeler, P. M., Barrett, L., Blumstein, D. T. & Clutton-Brock, T. H. Constraints and flexibility in mammalian social behaviour: introduction and synthesis. Philos T R Soc B 368 (2013). 10.1098/rstb.2012.0337

2 Komdeur, J. Variation in individual investment strategies among social animals. Ethology 112, 729–747 (2006). 10.1111/j.1439-0310.2006.01243.x

3 Hamilton, W. D. The genetical evolution of social behaviour. I. J Theor Biol 7, 1–16 (1964). 10.1016/0022-5193(64)90038-4

4 Hamilton, W. D. The genetical evolution of social behaviour. II. J Theor Biol 7, 17–52 (1964). 10.1016/0022-5193(64)90039-6

5 Eberhard, M. J. W. The evolution of social behavior by kin selection. The Quarterly Review of Biology 50, 1–33 (1975).

6 Page, A. E. et al. Testing adaptive hypotheses of alloparenting in Agta foragers. Nat Hum Behav 3, 1154–1163 (2019). 10.1038/s41562-019-0679-2

7 Perry, G. Going home: how mothers maintain natal family ties in a patrilocal society. Hum Nature-Int Bios 28, 219–230 (2017). 10.1007/s12110-016-9282-7

8 Briga, M., Pen, I. & Wright, J. Care for kin: within-group relatedness and allomaternal care are positively correlated and conserved throughout the mammalian phylogeny. Biol Letters 8, 533–536 (2012). 10.1098/rsbl.2012.0159

9 Lukas, D. & Clutton-Brock, T. Social complexity and kinship in animal societies. Ecol Lett 21, 1129–1134 (2018). 10.1111/ele.13079

10 Croft, D. P. et al. Kinship dynamics: patterns and consequences of changes in local relatedness. P Roy Soc B-Biol Sci 288 (2021). 10.1098/rspb.2021.1129

11 Ellis, S. et al. Patterns and consequences of age-linked change in local relatedness in animal societies. Nat Ecol Evol 6, 1766–1776 (2022). 10.1038/s41559-022-01872-2

12 Johnstone, R. A. & Cant, M. A. The evolution of menopause in cetaceans and humans: the role of demography. P Roy Soc B-Biol Sci 277, 3765–3771 (2010). 10.1098/rspb.2010.0988

13 Spong, G., Stone, J., Creel, S. & Björklund, M. Genetic structure of lions (Panthera leo L.) in the Selous Game Reserve: implications for the evolution of sociality. J Evolution Biol 15, 945–953 (2002). 10.1046/j.1420-9101.2002.00473.x

14 Dugdale, H. L., Macdonald, D. W., Pope, L. C., Johnson, P. J. & Burke, T. Reproductive skew and relatedness in social groups of European badgers,. Mol Ecol 17, 1815–1827 (2008). 10.1111/j.1365-294X.2008.03708.x

15 Croft, D. P. et al. Reproductive conflict and the evolution of menopause in killer whales. Curr Biol 27, 298–304 (2017). 10.1016/j.cub.2016.12.015

16 He, P. et al. Predicting kinship dynamics during pre-and post-reproductive life stages. bioRxiv (2025).

17 Buss, D. M. & Schmitt, D. P. Sexual strategies theory: an evolutionary perspective on human mating. Psychol Rev 100, 204–232 (1993). 10.1037/0033-295x.100.2.204

18 Buss, D. M. Conflict between the sexes: strategic interference and the evocation of anger and upset. J Pers Soc Psychol 56, 735–747 (1989). 10.1037/0022-3514.56.5.735

19 Fortunato, L. & Archetti, M. Evolution of monogamous marriage by maximization of inclusive fitness. J Evolution Biol 23, 149–156 (2010). 10.1111/j.1420-9101.2009.01884.x

20 Cant, M. A. & Johnstone, R. A. Reproductive conflict and the separation of reproductive generations in humans. P Natl Acad Sci USA 105, 5332–5336 (2008). 10.1073/pnas.0711911105

21 Hooper, P. L., Gurven, M., Winking, J. & Kaplan, H. S. Inclusive fitness and differential productivity across the life course determine intergenerational transfers in a small-scale human society. P Roy Soc B-Biol Sci 282 (2015). 10.1098/rspb.2014.2808

22 Goodenough, W. H. Residence rules. Southwest J Anthropol 12, 22–37 (1956). https://www.jstor.org/stable/3628856

23 Bergendorff, S. Kinship and human evolution: making culture, becoming human. (Lexington Books, 2016).

24 Fox, R. Kinship and marriage: an anthropological perspective. (cambridge university press, 1983).

25 Koster, J. et al. Kinship ties across the lifespan in human communities. Philos T R Soc B 374 (2019). 10.1098/rstb.2018.0069

26 Allen, N. J., Callan, H., Dunbar, R. & James, W. Early human kinship: from sex to social reproduction. (John Wiley & Sons, 2011).

27 Chen, Y., Ge, E. R., Zhou, L. Q., Du, J. & Mace, R. Sex inequality driven by dispersal. Curr Biol 33, 464–473 (2023). 10.1016/j.cub.2022.12.027

28 Henrich, J., Boyd, R. & Richerson, P. J. The puzzle of monogamous marriage. Philos T R Soc B 367, 657–669 (2012). 10.1098/rstb.2011.0290

29 Betzig, L. Causes of conjugal dissolution: a cross-cultural study. Curr Anthropol 30, 654–676 (1989). 10.1086/203798

30 Scelza, B. A. et al. High rate of extrapair paternity in a human population demonstrates diversity in human reproductive strategies. Sci Adv 6 (2020). 10.1126/sciadv.aay6195

31 Hughes, A. L. Evolution and human kinship. (Oxford University Press, 1988).

32 Ridgeway, C. L. Framed by gender: how gender inequality persists in the modern world. (Oxford University Press, 2011).

33 Stone, L. & King, D. E. Kinship and gender: an introduction. (Routledge, 2018).

34 Du, J. et al. Post-marital residence patterns and the timing of reproduction: evidence from a matrilineal society. P Roy Soc B-Biol Sci 290 (2023). 10.1098/rspb.2023.0159

35 UN. World Population Prospects 2022 (Department of Economic and Social Affairs, Population Division). (2024).

36 He, P. et al. Group size modulates kinship dynamics and selection on social traits. bioRxiv (2023).

37 Dyble, M., Migliano, A. B., Page, A. E. & Smith, D. Relatedness within and between Agta residential groups. Evol Hum Sci 3 (2021). 10.1017/ehs.2021.46

38 Wu, J. J. et al. Communal breeding promotes a matrilineal social system where husband and wife live apart. P Roy Soc B-Biol Sci 280 (2013). 10.1098/rspb.2013.0010

39 Richards, A. I. in African systems of kinship and marriage (Oxford University Press, 1950).

40 Schneider, D. M. Matrilineal kinship. (University of California Press, 1974).

41 Gaulin, S. J. C. & Schlegel, A. Paternal confidence and paternal investment: a cross-cultural test of a sociobiological hypothesis. Ethol Sociobiol 1, 301–309 (1980). 10.1016/0162-3095(80)90015-1

42 Fortunato, L. The evolution of matrilineal kinship organization. P Roy Soc B-Biol Sci 279, 4939–4945 (2012). 10.1098/rspb.2012.1926

43 Hartung, J. Paternity and inheritance of wealth. Nature 291, 652–654 (1981). 10.1038/291652a0

44 Eagly, A. H. & Wood, W. The origins of sex differences in human behavior: evolved dispositions versus social roles. American psychologist 54, 408 (1999). 10.1037/0003-066X.54.6.408

45 Archer, J. Sex differences in social behavior: are the social role and evolutionary explanations compatible? American Psychologist 51, 909 (1996). 10.1037/0003-066X.51.9.909

46 Wood, W. & Eagly, A. H. Biosocial construction of sex differences and similarities in behavior. Adv Exp Soc Psychol 46, 55–123 (2012). 10.1016/B978-0-12-394281-4.00002-7

47 Shennan, S. & Steele, J. The archaeology of human ancestry: power, sex and tradition. (Routledge, 2005).

48 Marlowe, F. W. Marital residence among foragers. Curr Anthropol 45, 277–284 (2004). 10.1086/382256

49 Wood, W. & Eagly, A. H. A cross-cultural analysis of the behavior of women and men: implications for the origins of sex differences. Psychol Bull 128, 699–727 (2002). 10.1037//0033-2909.128.5.699

50 Oota, H., Settheetham-Ishida, W., Tiwawech, D., Ishida, T. & Stoneking, M. Human mtDNA and Y-chromosome variation is correlated with matrilocal versus patrilocal residence. Nat Genet 29, 20–21 (2001). 10.1038/ng711

51 Ly, G. et al. Residence rule flexibility and descent groups dynamics shape uniparental genetic diversities in South East Asia. Am J Phys Anthropol 165, 480–491 (2018). 10.1002/ajpa.23374

52 Heyer, E., Chaix, R., Pavard, S. & Austerlitz, F. Sex-specific demographic behaviours that shape human genomic variation. Mol Ecol 21, 597–612 (2012). 10.1111/j.1365-294X.2011.05406.x

53 Demetrius, L. Adaptive value, entropy and survivorship curves. Nature 275, 213–214 (1978). 10.1038/275213a0

54 Johnstone, R. A. & Cant, M. A. Evolution of menopause. Curr Biol 29, R112–R115 (2019). 10.1016/j.cub.2018.12.048

