## APPENDIX for "Predicting the Patterns of Kinship Dynamics in Human Societies"

#### *Identifying the matrilocal and patrilocal societies in Koster et al*

We identify the typical matrilocal and patrilocal societies across the 19 datasets published by Koster et al[1], by testing whether the proportion of adult females is significantly different from that of adult males in each of these societies. When the test result is statistically significant for a given society, it is classified as matrilocal — if the proportion of adult females is higher than that of the adult males, or patrilocal — if the opposite is true (Table A1). When testing model predictions, we use the relatedness datasets from these identified societies.

**Table A1** The 2 matrilocal (PU and NI) and 7 patrilocal (MA, LA, CE, MS, GA, MP, AZ, TP, TA) societies identified from the published datasets by Koster et al[1] across 19 human societies.

| site | society | abbr. | % adult females | % adult males | p-value | type |
|---|---|---|---|---|---|---|
| 1 | Savannah Pumé | PU | 57.89 | 14.29 | 0.02 | MAT [*] |
| 2 | Miskito | MK | 53.33 | 21.43 | 0.60 | |
| 3 | Mayangna | NI | 80.65 | 38.89 | <0.01 | MAT [*] |
| 4 | Coastal Afro-Colombians | CA | 48.15 | 45.83 | 0.65 | |
| 5 | Inland Afro-Colombians | IA | 46.59 | 47.54 | 0.39 | |
| 6 | Dominica | DM | 94.63 | 96.6 | 0.70 | |
| 7 | Inland Emberá | IE | 60 | 61.54 | 1 | |
| 8 | Choyeros | MX | 73.68 | 81.4 | 0.22 | |
| 9 | The Gambia, 2 | GB | 80.5 | 89.39 | 0.05 | |
| 10 | Mosuo | MM | 82.14 | 91.36 | 0.21 | |
| 11 | Maya | MA | 84.15 | 94.74 | <0.01 | PAT [*] |
| 12 | Lamalera | LA | 83.76 | 95.43 | <0.01 | PAT [*] |
| 13 | Coastal Emberá | CE | 11.76 | 13.33 | 1 | |
| 14 | Maasai | MS | 25.93 | 31.25 | 0.68 | |
| 15 | The Gambia, 1 | GA | 70.23 | 86.42 | <0.01 | PAT [*] |
| 16 | Mosuo | MP | 62.69 | 88.14 | <0.01 | PAT [*] |
| 17 | Alakāpuram | AZ | 37.05 | 85.33 | <0.01 | PAT [*] |
| 18 | Tenpaṭṭi | TP | 30.57 | 91.84 | <0.01 | PAT [*] |
| 19 | Tanna | TA | 27.03 | 86.67 | <0.01 | PAT [*] |

#### *Relatedness datasets*

All the relatedness data used are published. Specifically, we used the relatedness data of individuals collected (**1**) from the bilocal *Agta* society published by Dyble et al.[2] (sex- and age-explicit), (**2**) from the matrilocal *Savannah Pumé* and *Mayangna*, the patrilocal *Maya*, *Lamalera*, *Gambia*, *Mosuo*, *Alakāpuram*, *Tenpaṭṭi* and *Tanna* societies published by Koster et al.[1] (sex- and age-explicit), and (**3**) from the duolocal *Mosuo* society published by Wu et al.[3] (sex- and age-group specific). To get the sex- and age-explicit estimates of the average relatedness for individuals in the duolocal Mosuo society[3], we fit the local polynomial regression models (i.e., the LOESS regression models) to the extracted age-group specific average relatedness values for each sex (with the WebPlotDigitizer[4]), and get the predicted average relatedness values at each available age (see details in the code supplemented).

***Supplementary figures***

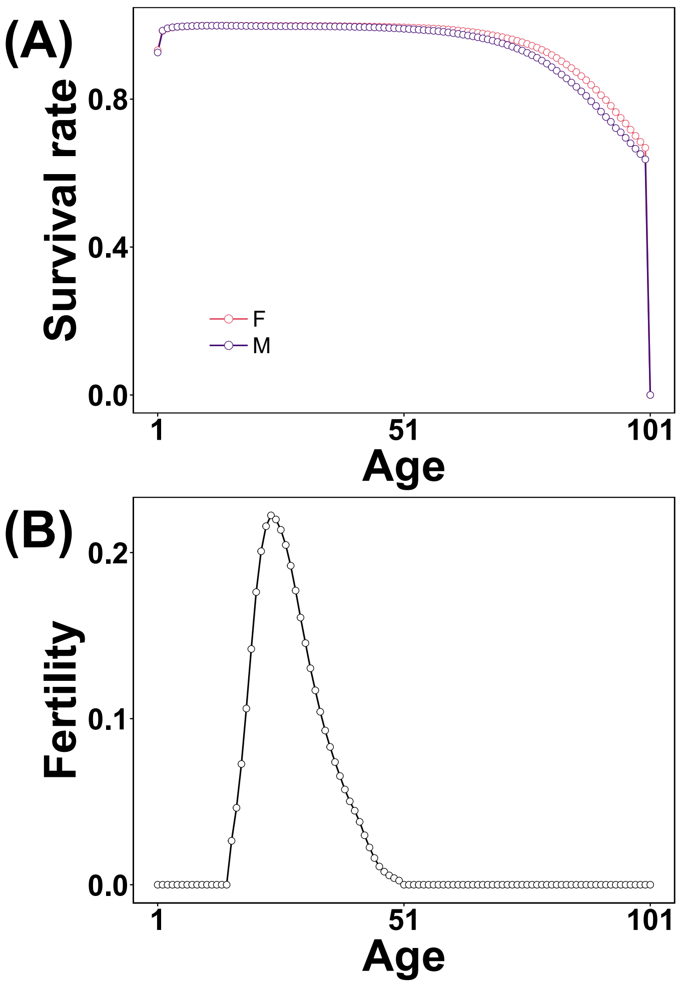


**Fig. A1** (**A**) The sex- and age-specific survival rates (derived from the 1986 survivorship data published by the UN[5]) are used for predicting the patterns of female and male kinship dynamics across the *bilocal* (BIL), *duolocal* (DUO), *matrilocal* (MAT) and *patrilocal* (PAT) societies. (**B**) The fertility schedule (derived from the 1986 global female fertility dataset published by the UN[5]) used for both sexes when predicting the patterns of human kinship dynamics across the 4 types of societies (due to the unavailability of male age-specific fertility data, we assume males have the same fertility schedule as females).

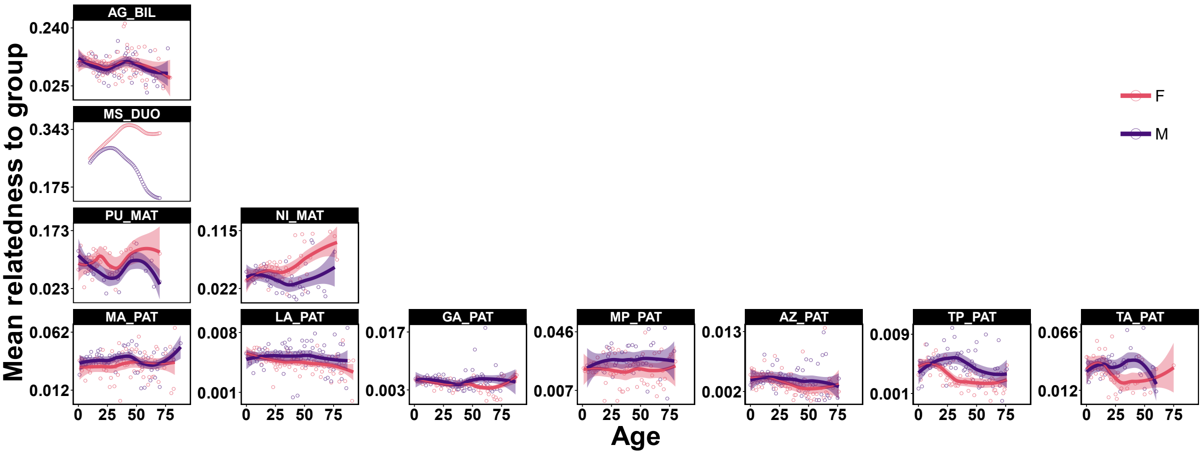


**Fig. A2** The observed patterns of sex- and age-specific mean relatedness of individuals to their groupmates (or household) in the *bilocal* (BIL), *duolocal* (DUO), *matrilocal* (MAT) and *patrilocal* (PAT) societies across the 11 human societies.

### References

1 Koster, J. *et al.* Kinship ties across the lifespan in human communities. *Philos T R Soc B* **374** (2019). https://doi.org/10.1098/rstb.2018.0069

2 Dyble, M., Migliano, A. B., Page, A. E. & Smith, D. Relatedness within and between Agta residential groups. *Evol Hum Sci* **3** (2021). https://doi.org/10.1017/ehs.2021.46

3 Wu, J. J. *et al.* Communal breeding promotes a matrilineal social system where husband and wife live apart. *P Roy Soc B-Biol Sci* **280** (2013). https://doi.org/10.1098/rspb.2013.0010

4 Rohatgi, A. WebPlotDigitizer. (2017).

5 UN. World Population Prospects 2022 (Department of Economic and Social Affairs, Population Division). (2024).
